# Do we really understand the role of the prefrontal cortex in placebo analgesia?

**DOI:** 10.1101/2021.06.18.449012

**Authors:** Eleni Frangos, Nicholas Madian, Binquan Wang, Megan L. Bradson, John L. Gracely, Emily A. Richards, Luana Colloca, Petra Schweinhardt, M. Catherine Bushnell, Marta Ceko

## Abstract

Several reviews have strongly implicated prefrontal cortical engagement in expectation-based placebo analgesia. We recently found a robust placebo analgesic response and associated decreases in pain-related cortical activations, without observable prefrontal engagement. We hypothesized our substantial conditioning and weak verbal instructions diminished expectation-related prefrontal activation. To test this, we examined the same subjects during a conditioning procedure, in which expectancy of pain relief was high. In two conditioning sessions, noxious heat was applied to a leg region treated with an “analgesic” cream and another treated with a “moisturizing” cream. In reality, both creams were inert, but the temperature applied to the moisturizing-cream area was 2°C higher than that applied to the analgesic-cream area.

Functional MRI was acquired during the second conditioning session. Pain ratings were lower for the low heat than the high heat, with corresponding reduced activations in pain-related regions. Similar to previous studies with strong expectation for pain relief, we observed more prefrontal activations during the “analgesic” than the control condition. Nevertheless, contrary to the idea of active prefrontal engagement, the relative activation was based on differences in *negative* BOLD signals. A literature review revealed that only a few studies conclusively showed active engagement of prefrontal cortex, i.e. increased positive BOLD signal during high expectation compared to a control, with variable timing and spatial-specificity. We suggest that this variability is due to the heterogeneous influence of cognitive, emotional and motivational factors. Future studies should attempt to unravel the multiple contributions to placebo responsiveness in the prefrontal cortex.

## INTRODUCTION

Placebo analgesia induction procedures comprise distinct elements which appear to have separable neurobiological underpinnings. For example, the effects of expectation-enhancing verbal instructions (e.g., “this ‘powerful analgesic’ will decrease your pain”) seem to be mediated by the opioid system, in that they can be blocked by naloxone [1], an opioid antagonist, and activate mu-opioid neurotransmission [51]. By contrast, the effects of associative learning-related conditioning, in which stimulus intensities are surreptitiously manipulated during a conditioning phase, may be mediated by other systems, such as the cannabinoid system [6].

Similarly, different regions of the brain may be involved in placebo analgesia depending on the method of induction. Typically, placebo induction paradigms include both expectation-inducing verbal instructions and conditioning through associative learning. Increased activation in prefrontal regions, such as the dorsolateral prefrontal cortex (DLPFC), ventromedial prefrontal cortex (VMPFC), and the rostral anterior cingulate cortex (rACC), have been widely implicated in the placebo effect by studies using paradigms that combine expectation-enhancing verbal instructions and conditioning [3; 18; 45]. However, in our recent placebo analgesia study [15], none of these regions were found to be activated during the placebo test phase in a large sample of healthy participants, despite the presence of a robust behavioral placebo effect. Additionally, the placebo effect induced in this study was not blocked by naloxone. We hypothesized that the lack of prefrontal activation or naloxone effect was due to the fact that in addition to the standard instructions regarding the effectiveness of the “analgesic” (placebo) cream (e.g., “this cream is a highly effective topical pain reliever”), the participants were also given certain instructions not typically given in most placebo studies (i.e., that the naloxone could block the effect of the “analgesic” [placebo] cream), which may have reduced the participants’ expectations of pain relief during the test phase. As a result, we proposed that the observed placebo effect was primarily driven by learning-related conditioning that did not activate opioidergic expectation-related prefrontal regions.

Importantly, fMRI data was also collected during the conditioning phase, prior to the test phase and the administration of naloxone. During this period, participants’ expectancy of pain relief should have been robust, as they had not yet been given the naloxone that they were told could block the effect of the “analgesic” (placebo) cream. Thus, the conditioning scan provided us with the opportunity to test our hypothesis, i.e., if the lack of prefrontal activation during the test phase was due to reduced expectancy caused by the drug administration, then prefrontal activations should be observed during the conditioning phase, prior to the drug administration, when expectancy of pain relief was high and reinforced by the conditioning trials. Here, we examined the anticipation and experience of pain relief during the conditioning scan in the same large sample of healthy participants studied in Frangos et al. [15] to determine whether regions of the prefrontal cortex (PFC) are engaged during high expectancy of pain relief.

## METHODS

### Participants

This study is part of a larger, previously published study that assessed placebo-induced analgesia in fibromyalgia patients compared to healthy controls [15]. The present study only includes 46 healthy participants (39 females, 7 males, mean age ± SD, 40 ± 13 years, range 19-64 years). The inclusion and exclusion criteria for the parent study are detailed in Frangos et al. [15]. In brief, the exclusion criteria for healthy participants included smoking of >10 cigarettes/week, alcohol consumption of >7 drinks/week for women and >14 drinks/week for men, use of recreational drugs and opioid medication, consumption of any pain medication other than NSAIDs within the past month or for more than one month on a continual basis within the past 6 months, pregnancy or breastfeeding, allergies to skin creams and lotions, chronic pain conditions, major medical, neurological, or current psychiatric conditions, including severe depression and generalized anxiety disorder, and MRI contraindications.

The study received approval from the NIH Institutional Review Board (IRB), and written informed consent was obtained from all participants according to the Declaration of Helsinki. As per IRB guidelines, the consent form included a general statement about deception: “At some point during the study we will give you misleading information. After the study is finished and all participants have been tested, we will explain how the information was not true and why.” No further details regarding deception were provided, and participants were not informed that the purpose of the study was to investigate placebo analgesia. Participants were compensated for completion of the study.

### Study design

The data presented here were collected during the second placebo conditioning phase (conditioning scan) of a larger placebo analgesia study [15]. The study included three placebo manipulation sessions (two sessions on day 1 in a mock scanner, and one session on day 2 during fMRI) and followed a well-established paradigm that included both verbally-induced expectation and conditioning components in a between- and within-subjects design [10; 13; 48].

The experimental design of the parent study is described in detail previously [15]. Briefly, participants were told that we were testing the mechanisms of a new powerful topical analgesic cream (the placebo cream) in comparison to a “hydrating” (control) cream. In actuality, both creams were identical. Participants came for testing on two separate days. On the first day, the “analgesic” cream was applied to two regions of the left leg and a “hydrating” cream to another two regions. After determining individual heat pain threshold and tolerance with a contact thermode, a mildly painful “low heat” stimulus was applied to one of the two “analgesic” regions, and a moderately painful “high heat” stimulus was applied to one of the two “hydrating” regions, with a temperature difference of ~2°C. On the second day, during the second placebo manipulation (conditioning scan), the same conditioning procedure was undertaken in the MRI scanner. After the conditioning scan, participants rated how effective they thought the cream was (0 = not effective at all, 10 = the most effective) and their desire for pain relief during the stimulus presentation (0 = no desire for pain relief, 10 = the most intense desire for pain relief imaginable). The placebo experimental scans subsequently followed and are detailed in Frangos et al., 2020.

#### Trial paradigm

The paradigm during the conditioning scan (Fig. 1) consisted of a baseline period (jittered 8-12 seconds; black crosshair on white background), an anticipation period (7 seconds; grayscale picture of control cream or placebo “analgesic” cream), a heat pulse (8.5 seconds; grayscale picture of thermode), a second anticipation period and heat pulse, a post-stimulus rest period (4 seconds), and 2 rating periods (7 seconds each for pain intensity [0 = no sensation, 100 = pain threshold, 200 = intolerable pain] and unpleasantness [-100 = extremely unpleasant, 0 = neutral, 100 = extremely pleasant]). Each heat pulse was presented on one of 2 pairs of treated 4 x 4 cm regions of the lower left leg.

**Figure 1.**
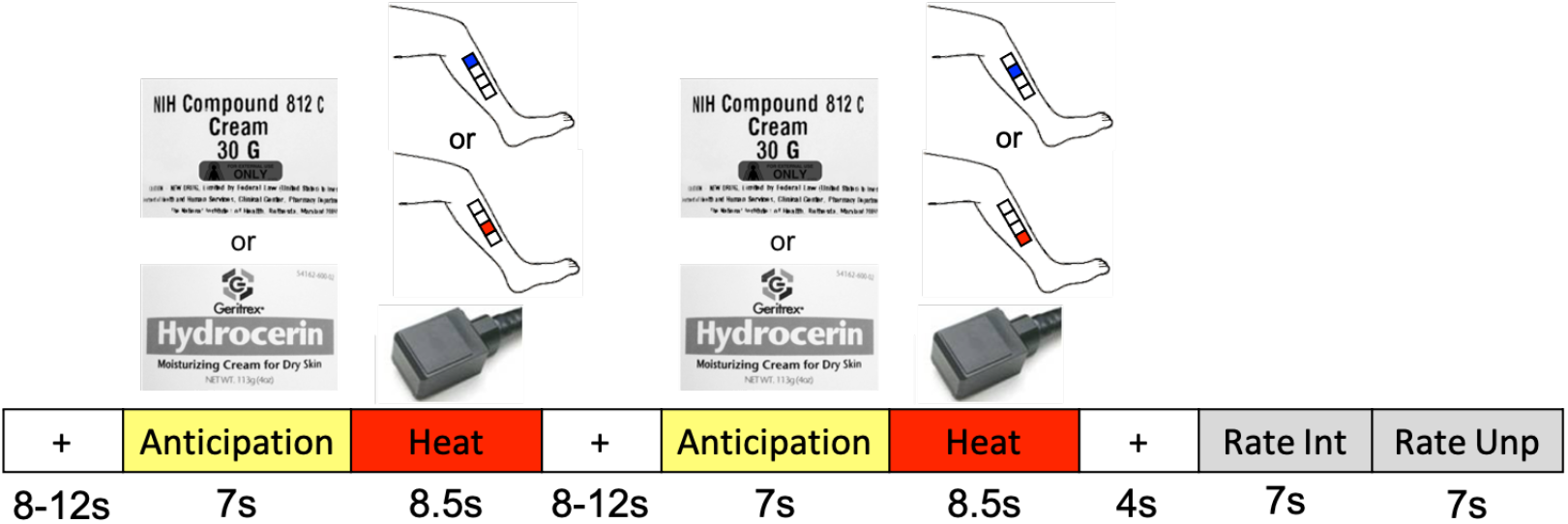
Experimental paradigm. The trial paradigm during the manipulation session consisted of a jittered interstimulus interval (ISI) of a black crosshair on white background, an anticipation period preceding the heat pulse (cue of either the “analgesic” (placebo) cream label (top image) or the “hydrating” (control) cream label (bottom image)), a low heat pulse (matched with the placebo cue) or a high heat pulse (matched with the control cue) on the left leg during which a thermode image was shown, a second jittered ISI, a second anticipation cue and matched heat pulse, a post-stimulus ISI, and a rating scale for pain intensity and pain unpleasantness. The blue and red squares on the leg images each indicate one of 2 pairs of 4 x 4 cm regions where the heat pulses were applied. *Adapted from Frangos et al., 2021*.

#### Behavioral data analysis

All behavioral measures were analyzed using 2-tailed paired t-tests or Wilcoxon signed-rank test for data that were not normally distributed based on the Shapiro-Wilk test. A significance level of p<0.05 was used in all analyses.

#### fMRI data pre-processing and analysis

Details of the fMRI acquisition and analysis methods are described in our previous publication [15]. In brief, each condition was modelled separately across trials (Fig. 1), i.e., the first anticipation period, first heat pulse, second anticipation period, second heat pulse and pain rating periods (intensity and unpleasantness combined) that occurred within a trial were each modelled as separate EVs for each condition (high heat [control cream] or low heat [placebo “analgesic” cream). For higher-level contrasts, voxel-wise thresholds were set to z > 3.1. If no differences were observed, the voxel-wise threshold was lowered to z > 2.3 to assess subtle effects and minimize false negatives (Type II error). All contrasts were cluster-corrected for multiple comparisons across the whole brain at p < 0.05.

## RESULTS

All 46 healthy participants were included in the analysis. A summary of demographic information can be found in the parent study [15]. Results are presented as mean ± SD.

### Manipulation Check: Low heat produced less pain and less neural activation than high heat

During the conditioning scan (second placebo manipulation), the high heat temperature administered on the “hydrating” (control) cream sites was 47°C ± 1.7°C, whereas the low heat temperature administered on the “analgesic” (placebo) cream sites was 44.6°C ± 1.8°C. As expected, the low heat condition was rated as less intense and unpleasant compared to high heat (intensity: high heat 151.7 ± 28.3, low heat 118.1 ± 34.3; unpleasantness: high heat 49.7 ± 23.9, low heat 20.5 ± 26.6; p’s <0.001). At the end of the conditioning scan, participants reported a moderate desire for pain relief (5.7 ± 3.1) and rated the “analgesic” (placebo) cream as moderately effective (5.5 ± 2.5).

The neural responses corroborated the perceptual responses as the low heat condition was associated with less activation in pain processing regions than the high heat condition. Although both high and low noxious heat stimuli produced activations within pain responsive regions (Table 1, left panel Fig. 2), the high heat produced significantly greater activation within regions that include the insula, secondary sensory cortices (SI and SII), and anterior cingulate cortex (ACC) (z > 3.1, p < 0.05, Table 1, right panel Fig. 2). The activation patterns and differences were consistently observed during both the first and second heat pulses.

**Table 1.**
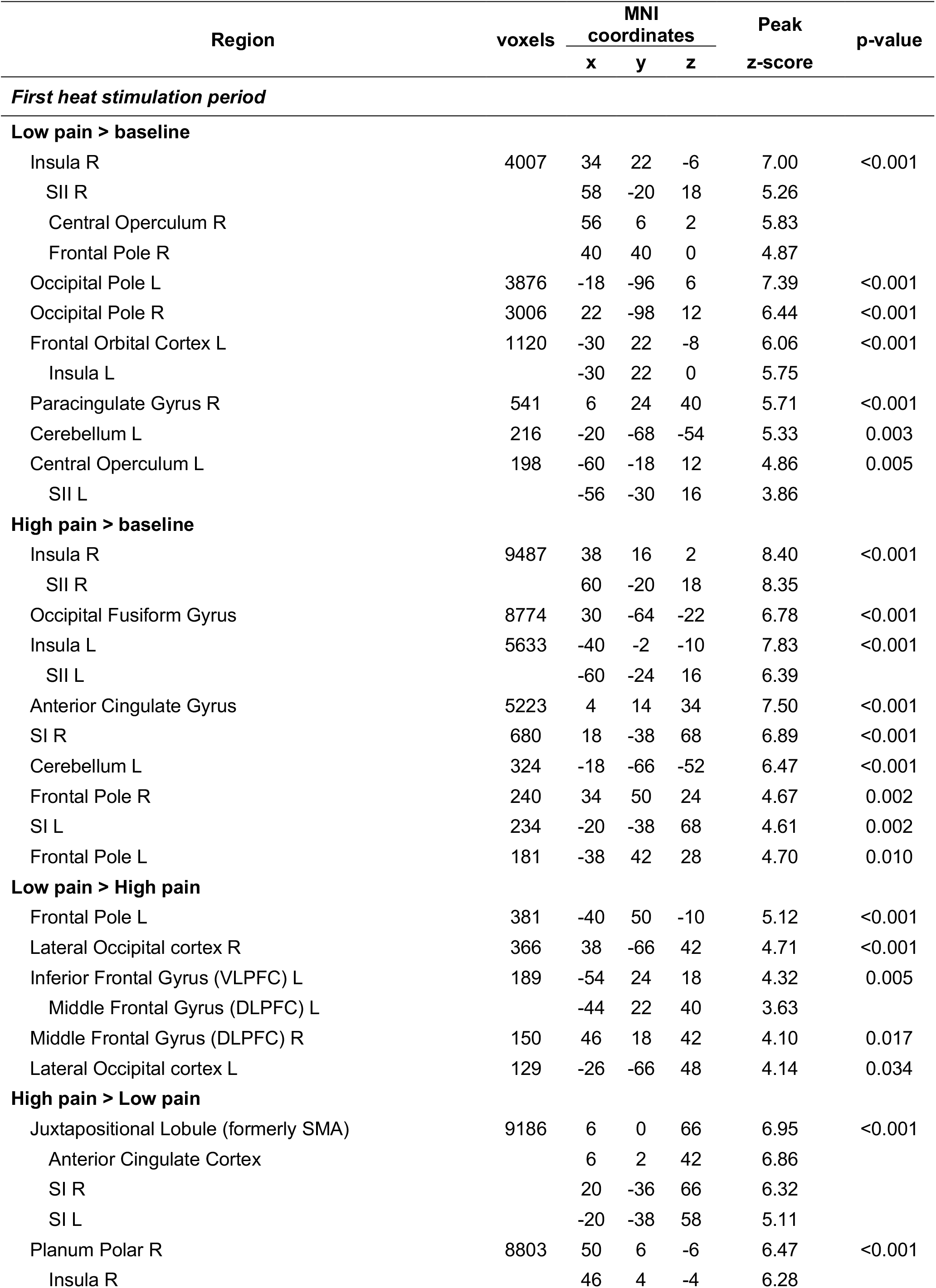
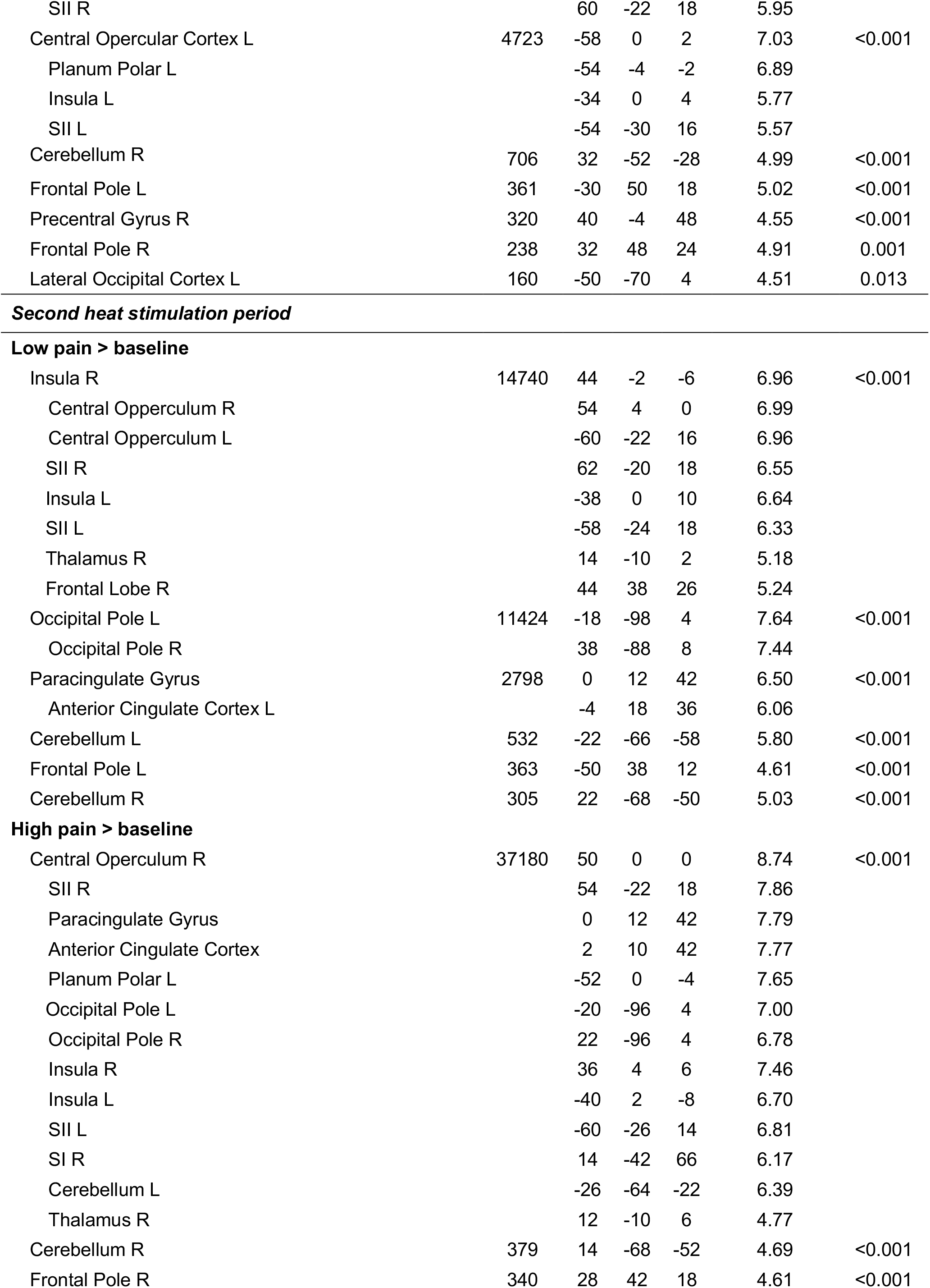
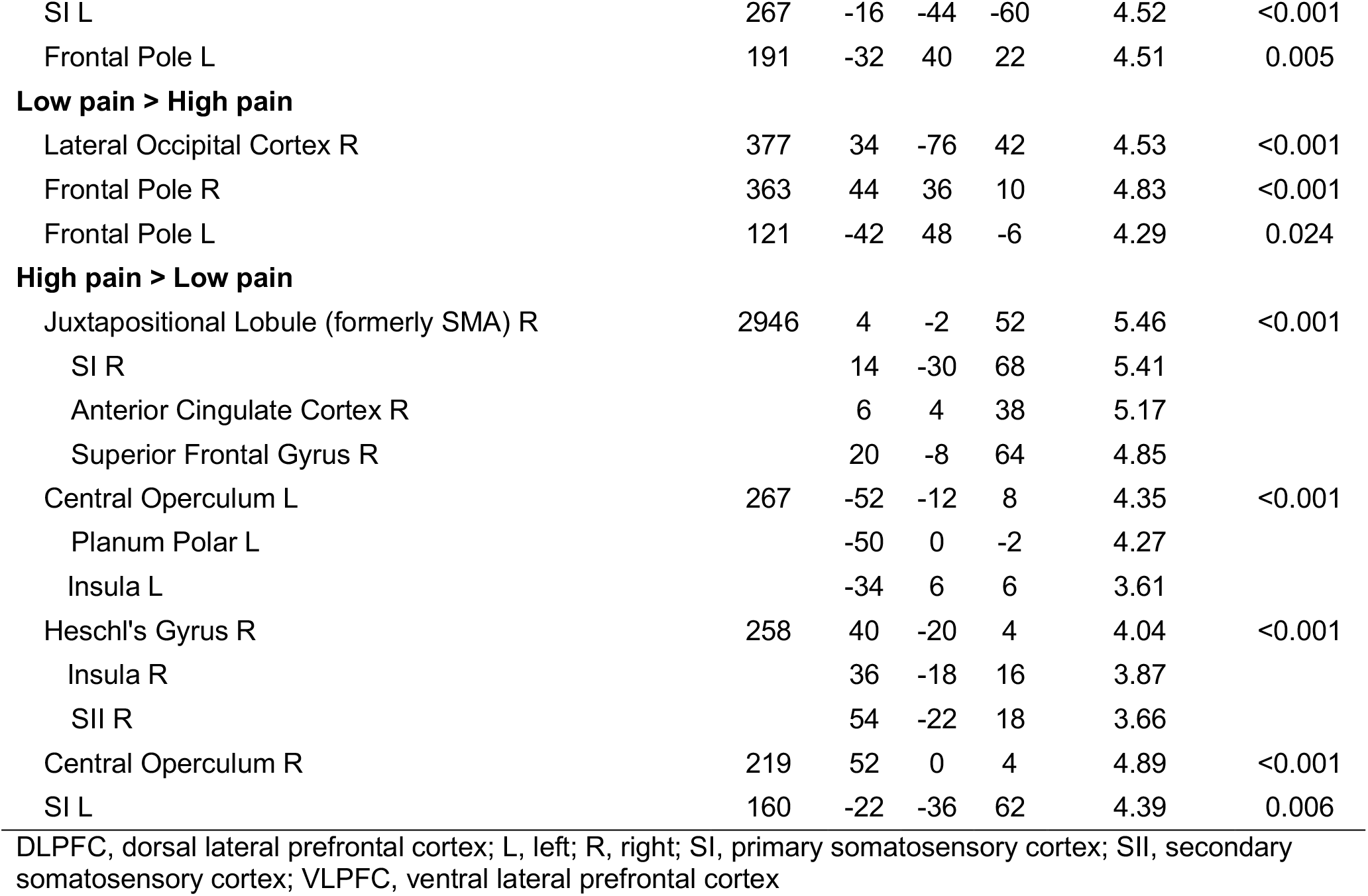
Whole-brain BOLD responses during the first and second heat stimulation periods.

| Region | voxels | MNI coordinates |  |  | Peak | p-value |
| --- | --- | --- | --- | --- | --- | --- |
|  |  | x | y | z | z-score |  |
| **First heat stimulation period** |  |  |  |  |  |  |
| **Low pain > baseline** |  |  |  |  |  |  |
| Insula R | 4007 | 34 | 22 | -6 | 7.00 | <0.001 |
| SII R |  | 58 | -20 | 18 | 5.26 |  |
| Central Operculum R |  | 56 | 6 | 2 | 5.83 |  |
| Frontal Pole R |  | 40 | 40 | 0 | 4.87 |  |
| Occipital Pole L | 3876 | -18 | -96 | 6 | 7.39 | <0.001 |
| Occipital Pole R | 3006 | 22 | -98 | 12 | 6.44 | <0.001 |
| Frontal Orbital Cortex L | 1120 | -30 | 22 | -8 | 6.06 | <0.001 |
| Insula L |  | -30 | 22 | 0 | 5.75 |  |
| Paracingulate Gyrus R | 541 | 6 | 24 | 40 | 5.71 | <0.001 |
| Cerebellum L | 216 | -20 | -68 | -54 | 5.33 | 0.003 |
| Central Operculum L | 198 | -60 | -18 | 12 | 4.86 | 0.005 |
| SII L |  | -56 | -30 | 16 | 3.86 |  |
| **High pain > baseline** |  |  |  |  |  |  |
| Insula R | 9487 | 38 | 16 | 2 | 8.40 | <0.001 |
| SII R |  | 60 | -20 | 18 | 8.35 |  |
| Occipital Fusiform Gyrus | 8774 | 30 | -64 | -22 | 6.78 | <0.001 |
| Insula L | 5633 | -40 | -2 | -10 | 7.83 | <0.001 |
| SII L |  | -60 | -24 | 16 | 6.39 |  |
| Anterior Cingulate Gyrus | 5223 | 4 | 14 | 34 | 7.50 | <0.001 |
| SI R | 680 | 18 | -38 | 68 | 6.89 | <0.001 |
| Cerebellum L | 324 | -18 | -66 | -52 | 6.47 | <0.001 |
| Frontal Pole R | 240 | 34 | 50 | 24 | 4.67 | 0.002 |
| SI L | 234 | -20 | -38 | 68 | 4.61 | 0.002 |
| Frontal Pole L | 181 | -38 | 42 | 28 | 4.70 | 0.010 |
| **Low pain > High pain** |  |  |  |  |  |  |
| Frontal Pole L | 381 | -40 | 50 | -10 | 5.12 | <0.001 |
| Lateral Occipital cortex R | 366 | 38 | -66 | 42 | 4.71 | <0.001 |
| Inferior Frontal Gyrus (VLPFC) L | 189 | -54 | 24 | 18 | 4.32 | 0.005 |
| Middle Frontal Gyrus (DLPFC) L |  | -44 | 22 | 40 | 3.63 |  |
| Middle Frontal Gyrus (DLPFC) R | 150 | 46 | 18 | 42 | 4.10 | 0.017 |
| Lateral Occipital cortex L | 129 | -26 | -66 | 48 | 4.14 | 0.034 |
| **High pain > Low pain** |  |  |  |  |  |  |
| Juxtapositional Lobule (formerly SMA) | 9186 | 6 | 0 | 66 | 6.95 | <0.001 |
| Anterior Cingulate Cortex |  | 6 | 2 | 42 | 6.86 |  |
| SI R |  | 20 | -36 | 66 | 6.32 |  |
| SI L |  | -20 | -38 | 58 | 5.11 |  |
| Planum Polar R | 8803 | 50 | 6 | -6 | 6.47 | <0.001 |
| Insula R |  | 46 | 4 | -4 | 6.28 |  |
| SII R |  | 60 | -22 | 18 | 5.95 |  |
| Central Opercular Cortex L | 4723 | -58 | 0 | 2 | 7.03 | <0.001 |
| Planum Polar L |  | -54 | -4 | -2 | 6.89 |  |
| Insula L |  | -34 | 0 | 4 | 5.77 |  |
| SII L |  | -54 | -30 | 16 | 5.57 |  |
| Cerebellum R | 706 | 32 | -52 | -28 | 4.99 | <0.001 |
| Frontal Pole L | 361 | -30 | 50 | 18 | 5.02 | <0.001 |
| Precentral Gyrus R | 320 | 40 | -4 | 48 | 4.55 | <0.001 |
| Frontal Pole R | 238 | 32 | 48 | 24 | 4.91 | 0.001 |
| Lateral Occipital Cortex L | 160 | -50 | -70 | 4 | 4.51 | 0.013 |
| **Low pain > baseline** |  |  |  |  |  |  |
| Insula R | 14740 | 44 | -2 | -6 | 6.96 | <0.001 |
| Central Opperculum R |  | 54 | 4 | 0 | 6.99 |  |
| Central Opperculum L |  | -60 | -22 | 16 | 6.96 |  |
| SII R |  | 62 | -20 | 18 | 6.55 |  |
| Insula L |  | -38 | 0 | 10 | 6.64 |  |
| SII L |  | -58 | -24 | 18 | 6.33 |  |
| Thalamus R |  | 14 | -10 | 2 | 5.18 |  |
| Frontal Lobe R |  | 44 | 38 | 26 | 5.24 |  |
| Occipital Pole L | 11424 | -18 | -98 | 4 | 7.64 | <0.001 |
| Occipital Pole R |  | 38 | -88 | 8 | 7.44 |  |
| Paracingulate Gyrus | 2798 | 0 | 12 | 42 | 6.50 | <0.001 |
| Anterior Cingulate Cortex L |  | -4 | 18 | 36 | 6.06 |  |
| Cerebellum L | 532 | -22 | -66 | -58 | 5.80 | <0.001 |
| Frontal Pole L | 363 | -50 | 38 | 12 | 4.61 | <0.001 |
| Cerebellum R | 305 | 22 | -68 | -50 | 5.03 | <0.001 |
| **High pain > baseline** |  |  |  |  |  |  |
| Central Operculum R | 37180 | 50 | 0 | 0 | 8.74 | <0.001 |
| SII R |  | 54 | -22 | 18 | 7.86 |  |
| Paracingulate Gyrus |  | 0 | 12 | 42 | 7.79 |  |
| Anterior Cingulate Cortex |  | 2 | 10 | 42 | 7.77 |  |
| Planum Polar L |  | -52 | 0 | -4 | 7.65 |  |
| Occipital Pole L |  | -20 | -96 | 4 | 7.00 |  |
| Occipital Pole R |  | 22 | -96 | 4 | 6.78 |  |
| Insula R |  | 36 | 4 | 6 | 7.46 |  |
| Insula L |  | -40 | 2 | -8 | 6.70 |  |
| SII L |  | -60 | -26 | 14 | 6.81 |  |
| SI R |  | 14 | -42 | 66 | 6.17 |  |
| Cerebellum L |  | -26 | -64 | -22 | 6.39 |  |
| Thalamus R |  | 12 | -10 | 6 | 4.77 |  |
| Cerebellum R | 379 | 14 | -68 | -52 | 4.69 | <0.001 |
| Frontal Pole R | 340 | 28 | 42 | 18 | 4.61 | <0.001 |
| SI L | 267 | -16 | -44 | -60 | 4.52 | <0.001 |
| Frontal Pole L | 191 | -32 | 40 | 22 | 4.51 | 0.005 |
| **Low pain > High pain** |  |  |  |  |  |  |
| Lateral Occipital Cortex R | 377 | 34 | -76 | 42 | 4.53 | <0.001 |
| Frontal Pole R | 363 | 44 | 36 | 10 | 4.83 | <0.001 |
| Frontal Pole L | 121 | -42 | 48 | -6 | 4.29 | 0.024 |
| **High pain > Low pain** |  |  |  |  |  |  |
| Juxtapositional Lobule (formerly SMA) R | 2946 | 4 | -2 | 52 | 5.46 | <0.001 |
| SI R |  | 14 | -30 | 68 | 5.41 |  |
| Anterior Cingulate Cortex R |  | 6 | 4 | 38 | 5.17 |  |
| Superior Frontal Gyrus R |  | 20 | -8 | 64 | 4.85 |  |
| Central Operculum L | 267 | -52 | -12 | 8 | 4.35 | <0.001 |
| Planum Polar L |  | -50 | 0 | -2 | 4.27 |  |
| Insula L |  | -34 | 6 | 6 | 3.61 |  |
| Heschl's Gyrus R | 258 | 40 | -20 | 4 | 4.04 | <0.001 |
| Insula R |  | 36 | -18 | 16 | 3.87 |  |
| SII R |  | 54 | -22 | 18 | 3.66 |  |
| Central Operculum R | 219 | 52 | 0 | 4 | 4.89 | <0.001 |
| SI L | 160 | -22 | -36 | 62 | 4.39 | 0.006 |

DLPFC, dorsal lateral prefrontal cortex; L, left; R, right; SI, primary somatosensory cortex; SII, secondary somatosensory cortex; VLPFC, ventral lateral prefrontal cortex

**Figure 2.**
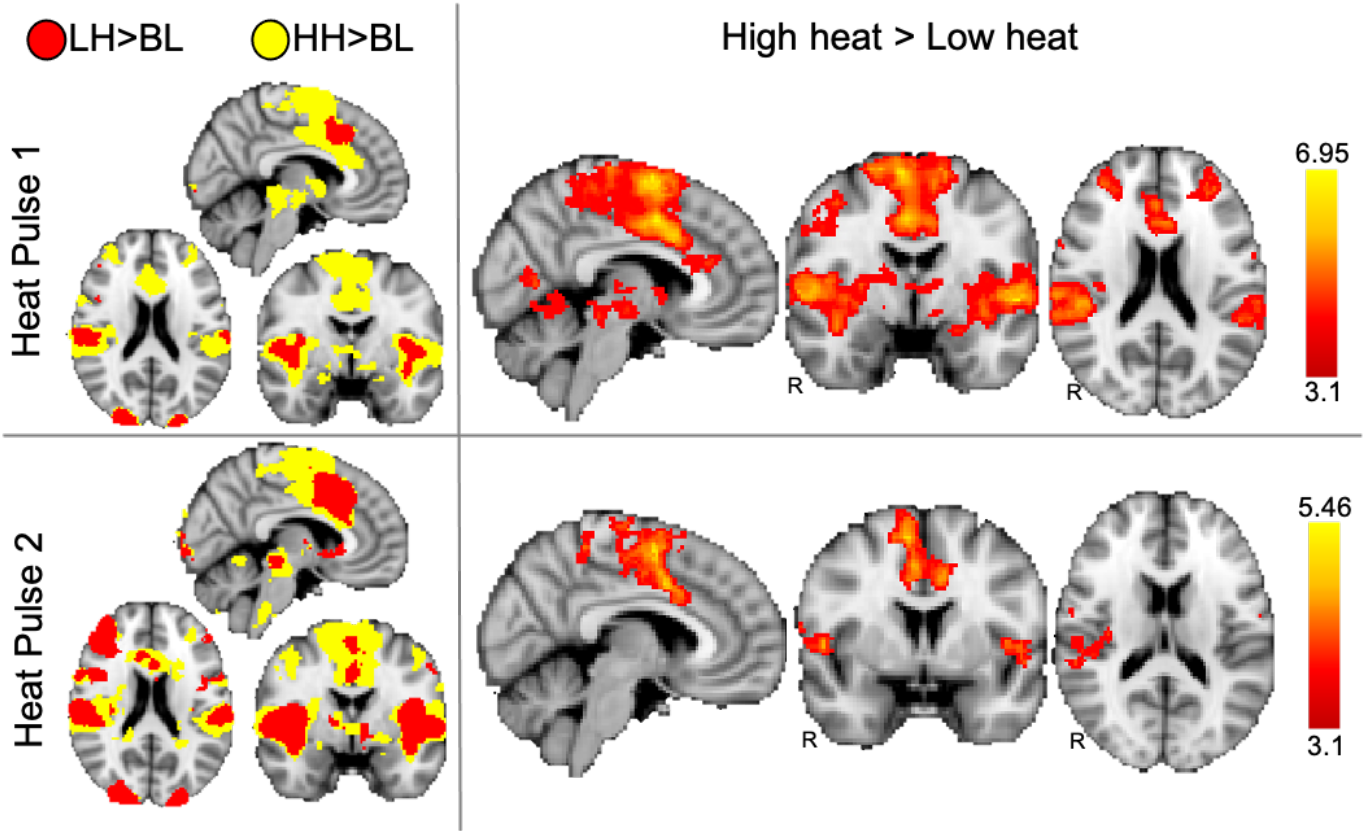
Neural responses to high and low painful heat. The left panel shows the pain-related activations in response to the low heat stimulus (red) and high heat stimulus (yellow) for each heat pulse. The right panel shows the difference in pain-evoked activations between the high and low heat stimuli for each heat pulse. All results are presented a voxel-based threshold of z > 3.1, cluster correction of p < 0.05. BL, baseline; HH, high heat; LH, low heat; R, right.

### Pain relief was associated with PFC activation resulting from a difference in negative BOLD signals

#### PFC activation during stimulation periods

In order to evaluate regions related to perceived pain relief, we examined regions that were activated more during low heat (when participants were feeling pain relief) than during high heat (no relief). Whereas high heat produced greater activations in pain-related regions as described above, during the first pulse, low heat produced greater activations in prefrontal regions, including DLPFC and VLPFC, as well as the lateral occipital cortex (z > 3.1, p < 0.05; Table 1; Figure 3A). Similarly, greater activity in the lateral occipital cortex and frontal pole were observed in the low heat > high heat contrast during the second pulse (z > 3.1, p < 0.05; Table 1; Figure 3B). Nevertheless, an examination of the parameter estimates (PEs, which provide BOLD signal directionality) extracted from the regions showing greater activation during low heat compared to high heat revealed that the observed differences between conditions during both the first and second stimulus pulses were based on differences in negative PE values, or deactivations, rather than positive PE values representative of activations above baseline (see graphs in Figure 3).

**Figure 3.**
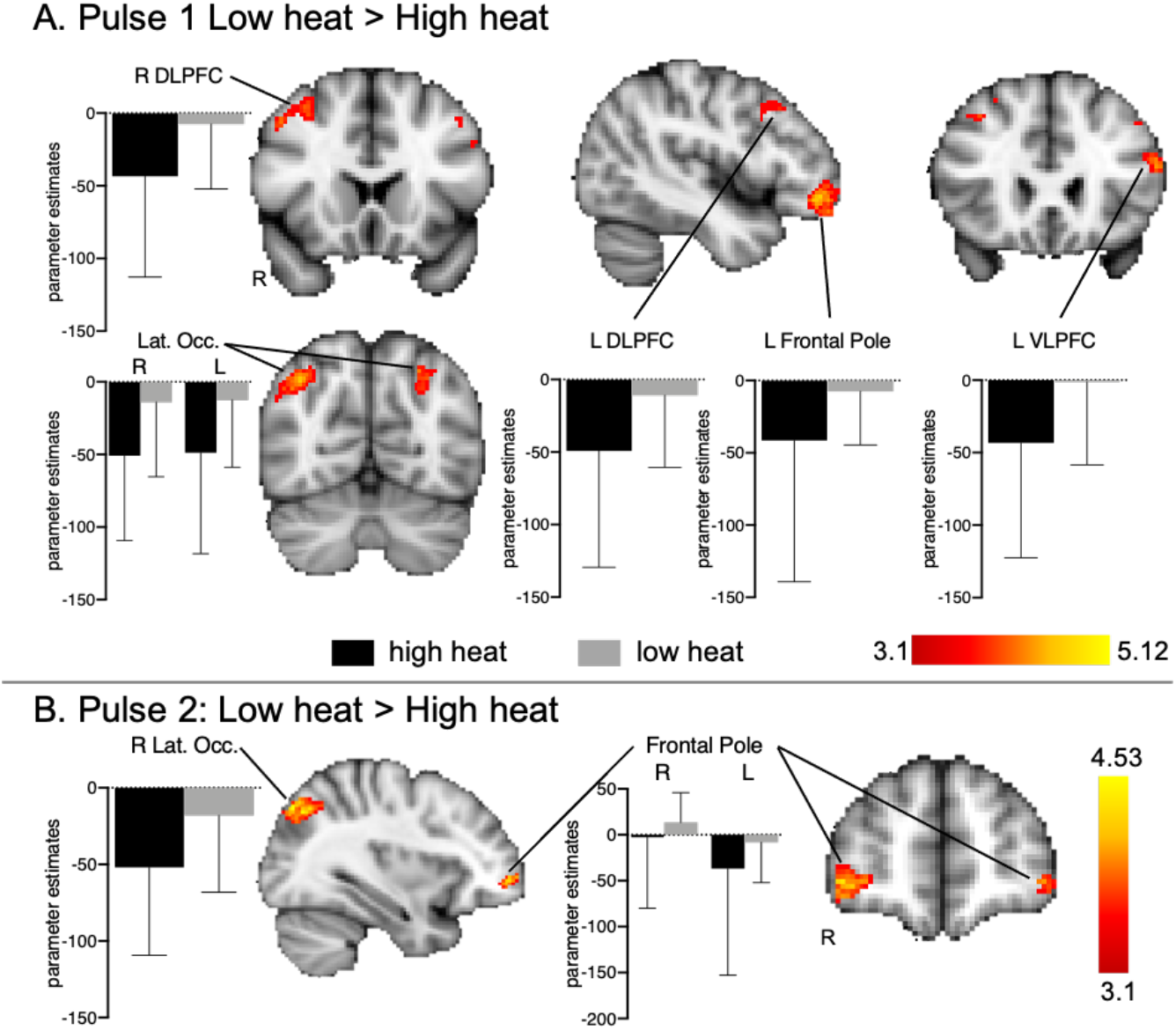
Neural responses during pain relief (low heat > high heat). Regions seemingly more activated during the first (A.) and second (B.) pulse of low heat are nearly all driven by negative parameter estimates, or *deactivations*, indicating that pain relief does not actively engage regions such as the DLPFC or VLPFC, as shown in the activation maps (voxel-based threshold z > 3.1, cluster correction of p < 0.05). L, left; R, right; DLPFC, dorsolateral prefrontal cortex; VLPFC, ventrolateral prefrontal cortex; Lat. Occ., lateral occipital cortex.

#### No PFC activation in anticipation periods

During the first and second heat pulse anticipation periods for both the high and low heat conditions compared to baseline, we found activations mainly within the visual cortex (z > 3.1, p < 0.05; left panel Fig. 4; Table 2), likely as a result of the visual cues presented during this period. Of particular interest was the low > high heat contrast for each anticipation period, as this contrast would indicate whether previously reported prefrontal networks are being engaged during anticipation of pain-relief. We did not observe any significant differences in either the first or second low pain anticipation > high pain anticipation contrasts, using a cluster forming threshold of z > 3.1 and cluster correction of p > 0.05. During anticipation of the second stimulus pulse, regions including the VLPFC and hippocampus were activated during anticipation of both high and low pain, but there was no difference between the conditions (Table 2, left panel Fig. 4). To examine more subtle effects and minimize false negatives, we decreased the cluster forming threshold to z > 2.3, and here we observed only minimal visual cortex activation in anticipation of low pain compared to high pain during the second stimulus pulse (right panel Fig. 4). No differences were observed in the high pain anticipation > low pain anticipation contrasts for both pairs of stimuli (Table 2).

**Figure 4.**
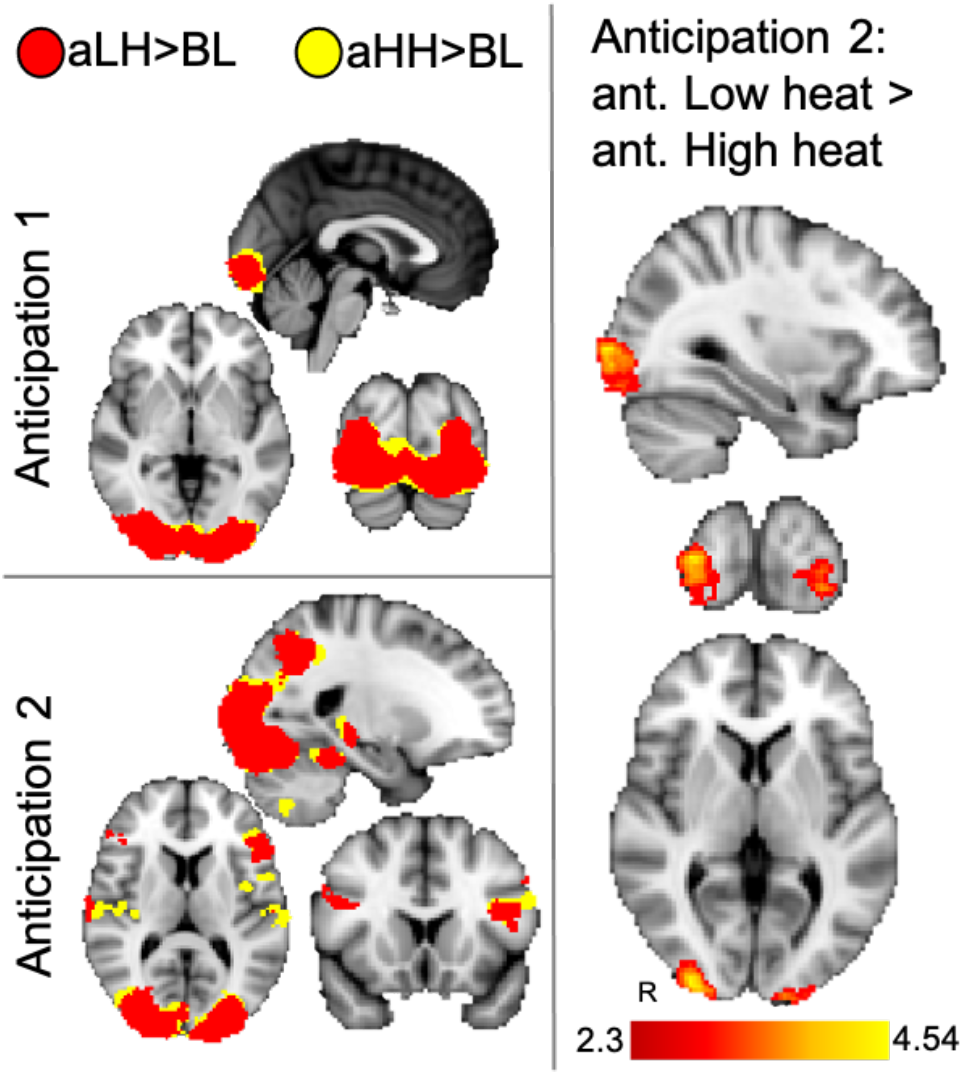
Anticipation-related activations. The left panel shows activations during the first and second period of anticipation (or expectation) of pain relief (low heat; red) and high heat (yellow) (voxel-based threshold z > 3.1, cluster correction of p < 0.05). The right panel shows significant differences within the occipital cortex in anticipation of low heat compared to high heat for only the second anticipation period, and only after decreasing the voxel-based threshold to z > 2.3 (cluster correction of p < 0.05). aHH, anticipation of high heat; aLH, anticipation of low heat; ant. anticipation; R, right.

**Table 2.** Whole-brain BOLD responses during the first and second anticipation periods.

| Region | voxels | MNI coordinates |  |  | Peak | p-value |
| --- | --- | --- | --- | --- | --- | --- |
|  |  | x | y | z | z-score |  |
| **First anticipation period** |  |  |  |  |  |  |
| **Low pain > baseline** |  |  |  |  |  |  |
| Occipital Pole L | 13641 | -16 | -98 | -2 | 9.14 | <0.001 |
| Occipital Pole R |  | 20 | -90 | -8 | 8.46 |  |
| Lateral Occipital Cortex L | 168 | -22 | -64 | 44 | 4.22 | 0.006 |
| **High pain > baseline** |  |  |  |  |  |  |
| Occipital Fusiform Gyrus R | 13701 | 18 | -90 | -10 | 8.69 | <0.001 |
| Occipital Pole |  | -16 | -100 | 0 | 8.54 |  |
| **Low pain > High pain** |  |  |  |  |  |  |
| - | - | - | - | - | - | - |
| **High pain > Low pain** |  |  |  |  |  |  |
| - | - | - | - | - | - | - |
| **Second anticipation period** |  |  |  |  |  |  |
| **Low pain > baseline** |  |  |  |  |  |  |
| Occipital Pole L | 20986 | -16 | -100 | 0 | 10.10 | <0.001 |
| Lateral Occipital Cortex R |  | 32 | -86 | -12 | 9.17 |  |
| Inferior Frontal Gyrus (VLPFC) L | 1186 | -40 | 14 | 18 | 5.24 | <0.001 |
| Precentral Gyrus L |  | -46 | -2 | 36 | 4.97 |  |
| Inferior Frontal Gyrus (VLPFC) R | 438 | 48 | 14 | 26 | 4.63 | <0.001 |
| Hippocampus L | 145 | -22 | -28 | -8 | 6.33 | 0.008 |
| Supramarginal Gyrus R | 130 | 68 | -20 | 18 | 3.93 | 0.014 |
| Hippocampus R | 108 | 22 | -28 | -8 | 6.13 | 0.034 |
| **High pain > baseline** |  |  |  |  |  |  |
| Occipital Pole R | 22333 | 12 | -90 | -8 | 8.81 | <0.001 |
| Occipital Pole L |  | -16 | -98 | 0 | 8.56 |  |
| Cerebellum R |  | 22 | -66 | -48 | 5.24 |  |
| S2 L |  | -60 | -24 | 18 | 4.63 |  |
| S1 R |  | 16 | -42 | 66 | 4.49 |  |
| Hippocampus R |  | 36 | -26 | -10 | 3.80 |  |
| Hippocampus L |  | -22 | -32 | -4 | 5.24 |  |
| Precentral Gyrus L | 1470 | -54 | 0 | 42 | 5.49 | <0.001 |
| Inferior Frontal Gyrus L (VLPFC) L |  | -42 | 8 | 22 | 5.46 |  |
| S1 R | 927 | 62 | -18 | 28 | 5.00 | <0.001 |
| Planum Temporale R |  | 64 | -18 | 12 | 5.00 |  |
| Planum Polar R |  | 48 | -4 | -2 | 4.42 |  |
| Insula R |  | 40 | 0 | -14 | 4.23 |  |
| S2 R |  | 36 | -30 | 22 | 4.11 |  |
| Cerebellum L | 161 | -22 | -68 | -52 | 4.66 | 0.005 |
| **Low pain > High pain** |  |  |  |  |  |  |
| *Occipital Pole R | 312 | 32 | -96 | 4 | 4.71 | <0.001 |
| *Occipital Pole L | 132 | -16 | -102 | 0 | 4.14 | <0.001 |
| **High pain > Low pain** |  |  |  |  |  |  |
| - | - | - | - | - | - | - |

\*, observed only in the analysis using cluster forming and cluster correction thresholds of $z > 2.3$ and $p < 0.05$ .

## DISCUSSION

Recently, we reported significant placebo analgesic responses associated with decreased pain-related cortical activations but failed to observe placebo-induced prefrontal activations [15]. We hypothesized that our paradigm, which involved substantial conditioning and weak verbal expectation instructions, reduced expectation-related prefrontal activation. Therefore, here, we examined the same healthy subjects during the conditioning session in which expectancy of pain relief was high, as expecations were not yet violated by the drug administration that would potentially block the effect of the “analgesic” (placebo) cream. As expected for a condition with less nociceptive input, the pain intensity and unpleasantness of the low heat were rated lower than the high heat and decreased activation was observed within pain responsive regions including insula, S2, S1, and ACC. Importantly, low heat pain compared to control high heat pain produced greater PFC activations, supporting our initial hypothesis and converging with the literature on expectation-induced placebo analgesia PFC responses. Since PFC effects were observed during the conditioning but not the placebo test phase [15], it is likely that the difference is due to lower expectancy during the test phase, suggesting that expectancy and conditioning effects may, indeed, have distinct neural pathways [6; 34].

### Engagement of the PFC during anticipation and experience of pain relief

Studies have shown that transient inhibition [26] and degeneration [7] of the PFC can block placebo analgesia. These findings, together with observations of differential PFC activation during placebo and control conditions [3], have been interpreted to indicate that control of subcortical regions via prefrontal engagement is necessary for expectation-based placebo analgesia [5]. Nevertheless, scrutinizing our own findings revealed an unexpected result: PFC “activations” observed during pain relief (low pain > high pain) were, in actuality, differences in negative BOLD responses, which seemed to contradict the prevalent theories. Thus, we questioned whether placebo-related PFC activations in other studies on healthy participants were also based on differences in negative BOLD signals. Table 3 summarizes the findings from studies included in two meta-analyses [2; 3] and more recent studies. Surprisingly, only six of the seventeen studies report PE values indicating BOLD signal directionality, which varied widely between and within studies. Taken together, these studies do not consistently corroborate the prevalent idea that expectation-based placebo is a result of increased PFC activity.

**Table 3.**
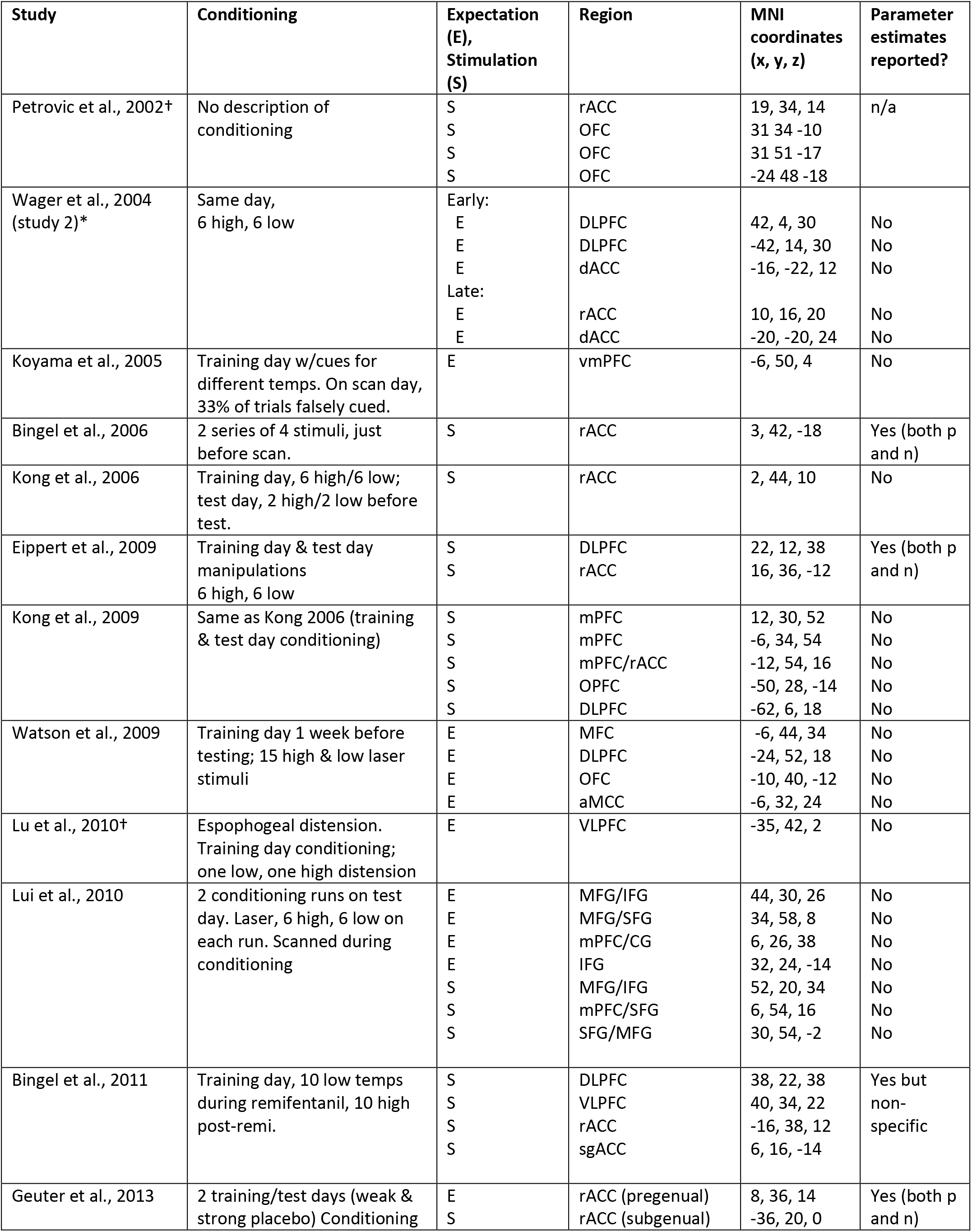
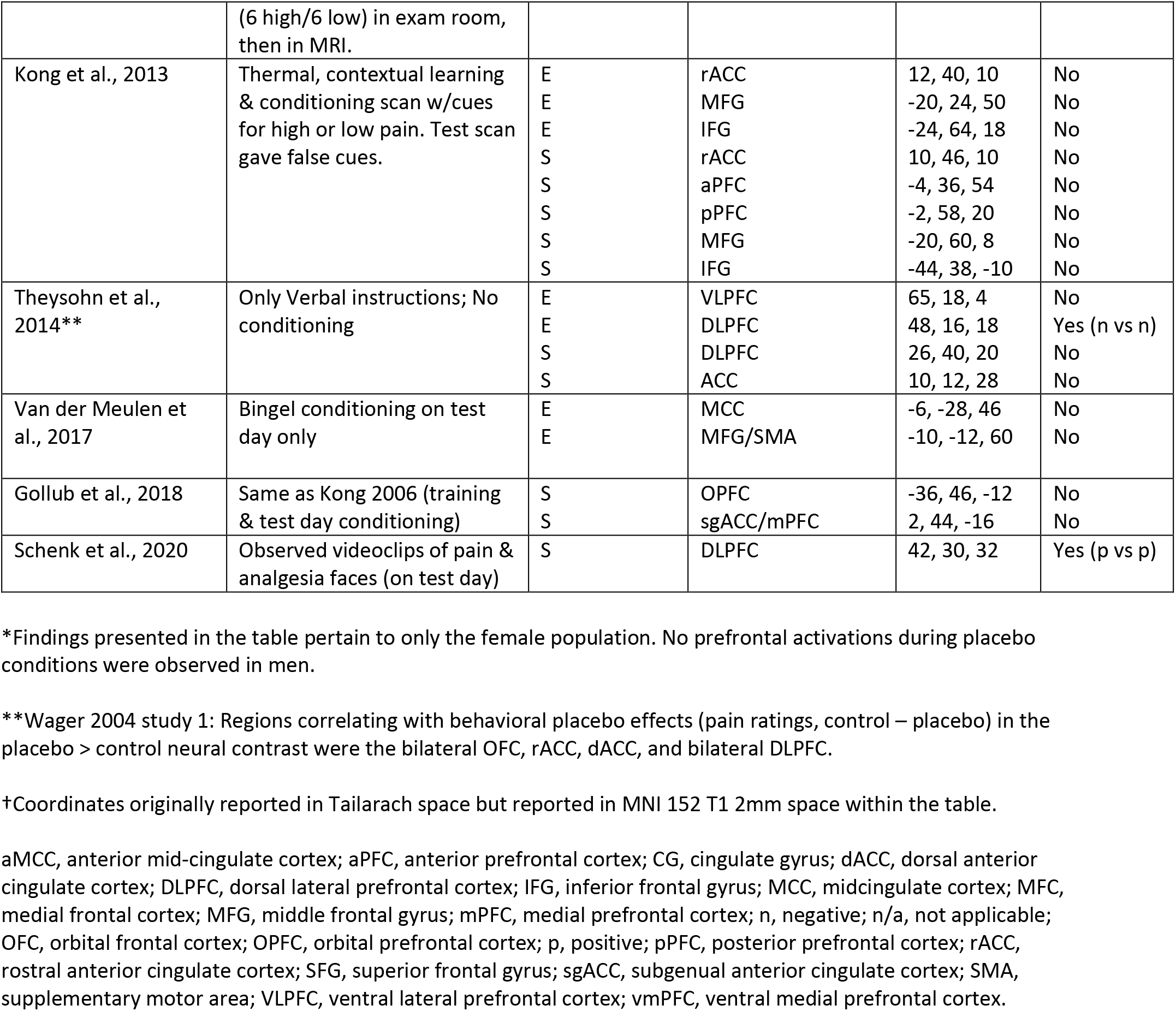
Frontal cortex activity during placebo analgesia expectation or stimulation periods in studies reporting on healthy participants.

PFC regions associated with placebo activity varied temporally and spatially across studies. Even the DLPFC and the rACC, the two regions in which placebo-related activity were most commonly observed, were identified only in seven [9; 13; 24; 35; 42; 48; 50] and eight [8; 9; 13; 17; 22; 23; 32; 48] of the seventeen studies, respectively. Other implicated PFC regions included the medial PFC (mPFC; four studies [19; 24; 30; 50]), orbitofrontal cortex (OFC; four studies [19; 24; 32; 50]), and ventrolateral prefrontal cortex (VLPFC; three studies [9; 29; 42]), among others [25; 43]. In fact, there was no single region where placebo-related activity was reported in a majority of the studies reviewed, and the lack of a discernable pattern among PFC regions associated with placebo analgesia during expectation or stimulation (Fig. 5), indicated that placebo analgesia involves parts of the PFC in some way, but there is great heterogeneity in how the PFC is implicated.

**Figure 5.**
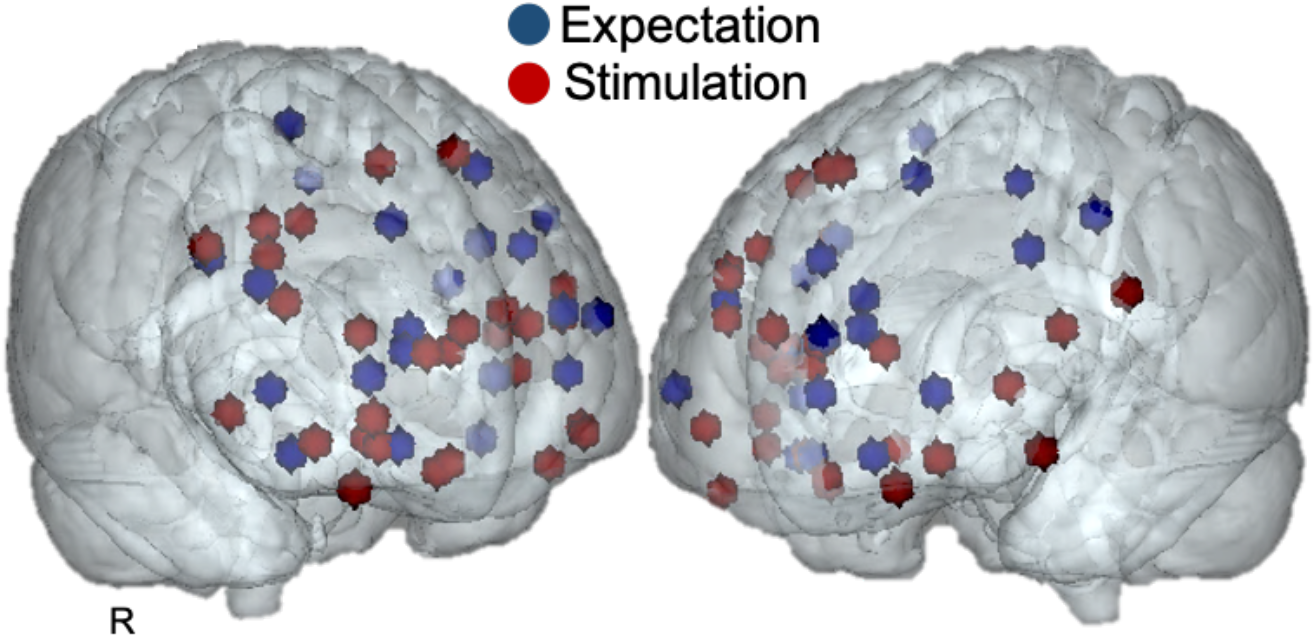
Localization of prefrontal cortex activations that have been reported during the expectation (blue) or stimulation (red) periods of placebo analgesia in studies conducted on healthy participants. Each sphere (rendered to 2mm on an MNI 152 T1 standard template) represents the peak prefrontal cortex coordinates reported in the studies (see Table 3 for details and references). R, right.

### Potential sources of PFC variability

The results of our study, in which the same subjects had differential PFC activation depending on the paradigm elements, support the conclusions of a recent meta-analysis by Zunhammer et al. [52]: that between-study variability in PFC activation corresponds to the heterogeneity of placebo induction paradigms. As an example, the divergent placebo effects in the pregenual rACC observed by Eippert et al. [13] and Geuter et al. [17] could be related to differences in expectations induced by their paradigms: Eippert et al. [13] administered saline or naloxone, whereas Geuter et al. [17] included two placebo efficacy conditions. However, differences in expectation are unlikely to be the only way design heterogeneity contributes to variability.

Other possible contributing factors are mood and attention, both of which modulate pain perception via different cortical circuitry [44]. Zunhammer et al. [53] suggest that different emotional regulation strategies, e.g., reappraisal or mindfulness, may be involved in placebo analgesia with differing levels of PFC activity. Yet, emotional regulation strategies may be influenced by setting-related [40] and cultural factors [11], which may each contribute differentially to between-study variability. The VMPFC plays a role in regulating negative affect [14], specifically in the extinction of the fear response to anticipated painful stimuli [36]. Thus, paradigms that induce negative affect, e.g., fear of painful stimuli, but also those that are conducive to the extinction of that fear, may induce greater VMPFC involvement. Other PFC responses to fear conditioning include DLPFC activations, and OFC and VMPFC deactivations [16]. Additionally, periaqueductal gray (PAG) activation has been observed during fear conditioning and early fear extinction [28]. These fear-related patterns of activity may be particularly prominent in studies that showed placebo-related activation during the anticipation period, since conditioned stimuli are used as cues.

Although more closely associated with subcortical areas, appetitive conditioning also involves the PFC. Exposure to appetitive-conditioned stimuli activates the medial OFC [21; 27], whereas inhibition of conditioning reinstatement activates the VMPFC [12]. As relief from pain is rewarding [4; 31], it is not surprising that these brain regions are engaged during placebo analgesia, but PFC engagement may vary depending on the induction and test paradigms, particularly timing factors. For example, OFC activation is associated with the acquisition of appetitive conditioning, but not with early or late extinction, whereas rACC activation is associated with the early extinction of appetitive conditioning, but not acquisition or late extinction [27]. Consequently, the variability observed within these regions may reflect the extent, duration, and efficacy of the placebo conditioning. For instance, the present neural activity was observed during a second conditioning session, which may have been too late to observe OFC involvement. Importantly, the rACC activation frequently observed in placebo studies may partially reflect the effects of early extinction, as most studies examine the placebo test phase when the placebo and control stimuli are equalized. Moreover, previous reports of rACC--PAG coupling during placebo analgesia [8; 13; 32; 49] may not only be reflective of pain modulation but also the moment of simultaneous extinction of appetitive and fear conditioning. Appetitive conditioning-induced activations are also influenced by personality traits, e.g., neuroticism [21] and sensation-seeking [41], that are dopamine-associated and correlate positively with placebo responsiveness [37], suggesting an intrinsic relationship among appetitive conditioning, dopaminergic neurotransmission, personality, and the placebo effect.

With regard to the role of attentional processes in placebo analgesia, related DLPFC activation has been suggested to reflect the redirection of attention away from the stimulus or *toward* the stimulus to evaluate treatment efficacy [48]. Thus, one might expect DLPFC activity in any situation in which participants are induced to direct attention toward or away from a stimulus, which is likely to occur repeatedly during placebo studies. DLPFC activation or deactivation might also reflect the degree of engagement of attentional and task-related networks. When painful stimuli are presented during a cognitive task, activity in the dorsomedial DLPFC decreases, while activity in the ventrolateral DLPFC increases [38]. Presumably, this corresponds respectively to deactivation of the default mode network [39] and activation of the extrinsic mode network [20], a generalized network involved in allocating cognitive resources to sensory processing or task performance. Thus, the level of attentional engagement in the paradigm may determine whether, where, and in what direction DLPFC activity occurs. Similarly, distractors may engage executive functions such as inhibition and set shifting, producing variable DLPFC and VLPFC activity across studies [33; 46; 47].

### Study limitations and suggestions for future placebo analgesia studies

In the present study, the psychological and cognitive variables described above were not measured and controlled for. These factors should be considered in future studies to obtain a clear understanding of the role of the PFC in placebo analgesia. Furthermore, clarity and standardization of instructions could improve the control of directed attention and expectation.

Also, when considering the early and late stages of pain in placebo studies, we should be mindful of the early and late contributions of conditioning and extinction. Finally, consistent reporting of the control and placebo conditions versus baseline to understand BOLD signal directionality is critical for interpreting subsequent contrasts used to assess functionality.

In conclusion, our study confirms previous findings of PFC activity during pain relief, relative to a control condition, when expectancy of pain relief is strong. Nevertheless, the relative activation is based on differences in negative BOLD signals contrary to the prevalent theory that active engagement of PFC is the underlying mechanism for expectation-related placebo analgesia. Further, examination of the literature revealed that only a few studies show active PFC-engagement during placebo analgesia, and the timing and spatial-specificity of activity varies widely likely due to uncontrolled psychological and cognitive factors. Therefore, to better understand the role of the PFC in placebo analgesia, future studies should control for various psychological and cognitive factors, and shift from asking *is the PFC involved* to *when, how* and *which sub-regions are involved*.

## Acknowledgements

This research was supported by the Intramural Research Program of the NIH, National Center for Complementary and Integrative Health. The authors thank Brian Walitt, Nicole Godwin, Linda Ellison-Dejewski, Brenda Justement, Susan Goo, and Patrick Korb for subject recruitment and clinical support.

